# Slower, but deeper community change: anthropogenic impacts on species temporal turnover are regulated by intrinsic dynamics

**DOI:** 10.1101/2022.11.10.515930

**Authors:** J. Christopher D. Terry, Axel G. Rossberg

## Abstract

Understanding the mechanisms behind biodiversity is central to assessing and forecasting anthropogenic impacts on ecological communities. However, quite how intrinsic ecological processes and external environmental drivers act together in natural systems to influence local temporal turnover is currently largely unexplored. Here, we determine how human impacts affect multiple metrics of bird community turnover to establish the ecological mechanisms behind compositional change. We used US Breeding Bird Survey data to calculate transect-level rates of three measures of temporal species turnover: a) short-term (initial rate of decline of Sørensen similarity), b) long-term (asymptotic Sørensen similarity), and c) overall species accumulation rate (species-time relationship exponents) over 2692 transects across 27 habitat types. We then hierarchically fit linear models to estimate the effect on these turnover metrics of anthropogenic impact via the Human Modification Index proxy, while accounting for observed species richness, the size of the species pool and annual environmental variability. We found broadly consistent impacts of increased anthropogenic pressures across diverse habitat types. The Human Modification Index was associated with greater turnover at long-timescales, but marginally slower short-term turnover. The species accumulation rate through time was not notably influenced. Examining anthropogenic impacts on different aspects of species turnover in combination allows greater ecological insight. Observed human impacts on short-term turnover were the opposite of existing expectations and suggest humans are disrupting the background turnover of these systems, rather than simply driving rapid directed turnover. The increased long-term turnover was driven by more frequent species changes between core and occasional status rather than greater arrival of ‘new’ species. These results highlight the role of intrinsic dynamics and caution against simple interpretations of increased species turnover as reflections of environmental change.

**Open Research Statement:** No new empirical data are used in the manuscript as all primary data is publicly available, as cited in the manuscript. Our code repository (including fitted model objects and markdown documents detailing all steps) is available at https://figshare.com/s/f5b9152ff7643efb347d.

## Introduction

Ecological communities are in a continual state of flux as they respond to changing external conditions and their internal dynamics play out. Understanding the drivers of this change in ecological communities is central to both core theories of biodiversity (MacArthur and Wilson 1967, Rosenzweig 1995, Hubbell 2001) and applied efforts to link biodiversity change to anthropogenic pressures (Magurran 2016). Even where tallies of species richness (α- diversity) through time are relatively constant, there is often considerable turnover in species composition (temporal *β*-diversity (Dornelas et al. 2014) and this turnover appears to be considerably faster than would be expected by neutral models (Gotelli et al. 2017). However, quantifying and summarising these changes remains a challenge as *β*-diversity is a fundamentally multi-facetted concept (Anderson et al. 2011). Since species richness trends are not necessarily correlated with wider biodiversity trends the benefits of simultaneously considering a broader spectrum of ecological responses to anthropogenic pressures are becoming increasingly clear (Hillebrand et al. 2018, Magurran et al. 2018).

Despite its acknowledged importance, temporal *β*-diversity has received less attention than spatial patterns, at least in part due to the challenges in gathering sufficient data (Magurran et al. 2019). Meta-analyses of observations of year-on-year turnover rates have identified latitudinal gradients in turnover (Korhonen et al. 2010), higher rates of change in marine systems (Blowes et al. 2019), identified a link between temperature changes and turnover rates in marine systems (Antão et al. 2020), and interactions between ‘naturalness’ and the rate of climate change (Pilotto et al. 2020) on turnover. Community change is frequently interpreted as a negative consequence or signal of human activity. However, fundamental ecological questions about the mechanisms driving change remain to be comprehensively addressed (Dornelas et al. 2023) which limit our capacity to interpret observations.

The similarity of a community as recorded between two timepoints is determined by a multitude of processes. Leaving sampling effects to one side for now, fundamentally these can be categorised into intrinsic and extrinsic drivers (Fig 1). Intrinsic turnover incorporates the processes that would generate ongoing turnover in local communities without long-term external change. This includes the impact of demographic stochasticity, dynamics generated by interspecific interactions between species within the existing metacommunity (O’Sullivan et al. 2021) and any local species turnover driven by ‘background’ environmental fluctuations (Kalyuzhny et al. 2014). By contrast, extrinsic drivers change the fundamentals of the community, and include long term climate and land-use changes and shifts in the species pool in the wider metacommunity (whether through range shifts or longer-term processes such as speciation).

**Fig 1.**
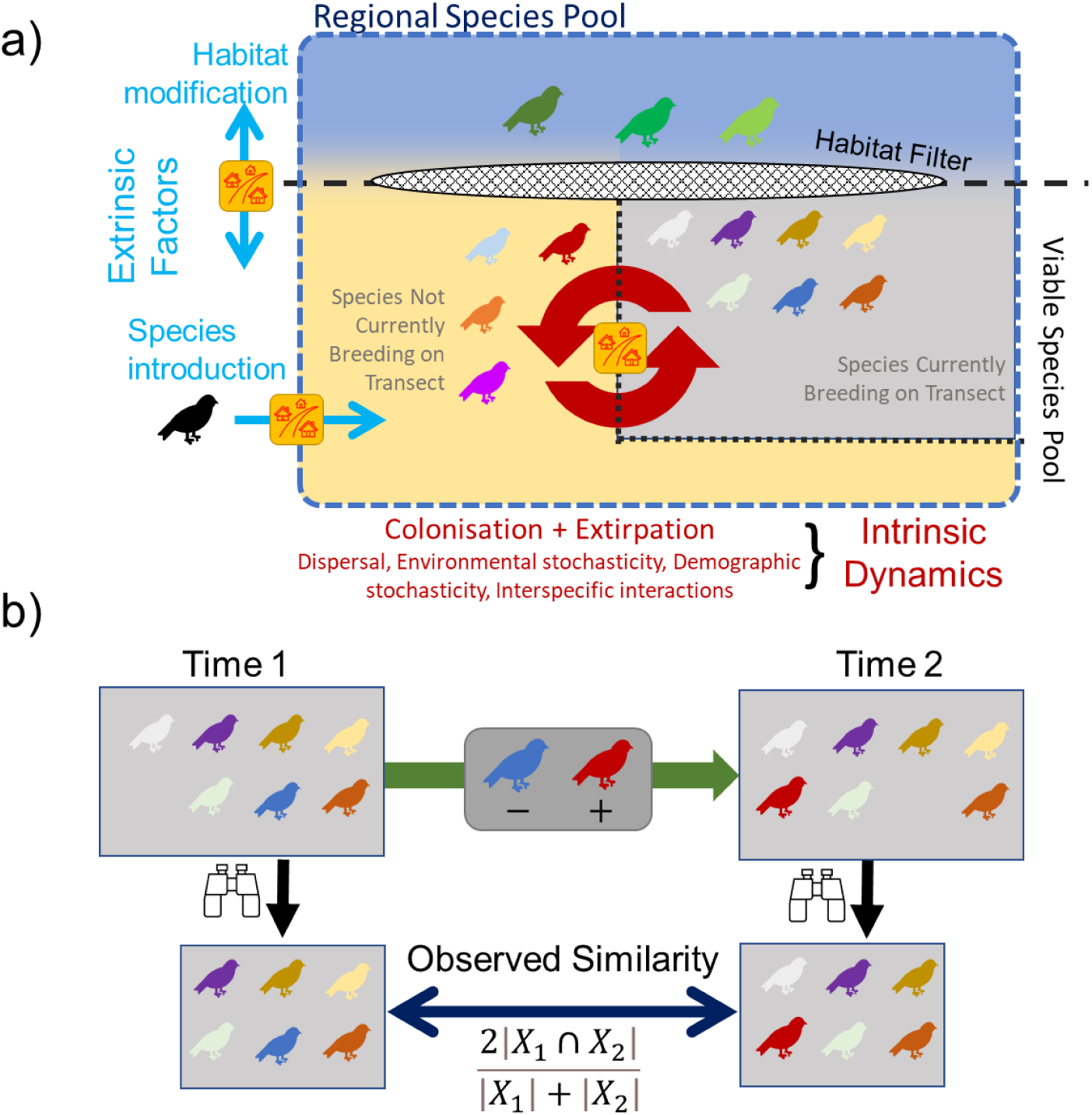
a) Cartoon depicting how both extrinsic factors and intrinsic dynamics both contribute to the species that are currently resident on a particular transect. Extrinsic habitat conditions set the overall filter controlling the viable species pool from the regional species pool, but only some of these viable species will actually be present at the local scale due to ongoing dynamics. Processes intrinsic to the system lead to a continual turnover in species that are currently resident at the focal site (red arrows). Species introductions can add to the regional species pool while human impacts (represented by the yellow house icon) could affect the majority of these processes. b) The observed similarity of ecological communities between two time points can be affected by both intrinsic and extrinsic drivers, making disentangling the underlying drivers a significant challenge. Here, the blue bird is lost and the dark red bird is gained between the observation points. The observed similarity will be deflated due to observation errors (principally false-absences).

Earlier work with island bird communities (Russell et al. 1995) suggested that the timescales at which these groups of processes were dominant were sufficiently different that they could be differentiated. However, given the rate at which global change is progressing, this assumption is unsafe. Extrinsic changes may plausibly be occurring on similar ‘intermediate’ time scales to intrinsic turnover and extensive previous work on North American birds has demonstrated that some bird species’ range and abundance changes are attributable to extrinsic drivers (Rosenberg et al. 2019, Di Cecco and Hurlbert 2022). Better differentiating the extent to which observed species turnover through time at a particular location is determined by external changes to the ecosystem or represents dynamics intrinsic to a given system would greatly aid the attribution of changes to particular drivers (Gonzalez et al. 2023).

The two aspects of temporal *β*-diversity most frequently studied to date have been: 1) the slopes of species accumulation curves (also termed the species-time relationship, STR (White et al. 2006, Rivett et al. 2021) and 2) measures of the rates of species gains and losses (Blowes et al. 2019, Antão et al. 2020, Pilotto et al. 2020). The second can either be directly reported or captured by similarity measures such as the Sørensen or Jaccard indices, which can then be partitioned to account for changing species richness (Baselga 2010, Tatsumi et al. 2021). Both aspects have been referred to as ‘turnover’, although they capture distinct ecological patterns. In particular, the STR captures the rate of appearance of ‘new’ species, while composition similarity measures could indicate high turnover where the same species repeatedly colonised and are then extirpated. The exponent of the STR is affected by the dynamics of rare or ephemeral species, but does not capture information about the permanence or otherwise of persistent species that are frequently observed.

Nevertheless, any single rate of ‘turnover’ is inevitably an average over the community, obscuring the inter-specific differences in residence time, extinction and recolonisation probability (Gotelli et al. 2022) and differentiation between ‘core’ and ‘occasional’ species (Magurran and Henderson 2003). Unless there are strong cyclic dynamics (Collins et al. 2000), the similarity of communities will decline through time. However, across ‘ecological’ time scales in non-successional systems we would expect the decline in similarity to asymptote at a value considerably above zero (Russell et al. 1995). Determining this asymptote requires long time series, but can be highly informative about the nature of the underlying ecological dynamics. In particular and in contrast to short-term turnover, the asymptote describes the ‘depth’ of the turnover and is influenced by the proportion of species that can be considered core species – those abundant and persistent species strongly associated with the habitat (Magurran and Henderson 2003).

Here we take a multiview perspective of turnover to examine the fundamental drivers of ecological change in a uniquely large and methodologically consistent dataset, the North American Breeding Bird Survey (BBS, Pardieck et al. 2020). We quantify a suite of aspects of species turnover for transect routes that met our criteria for inclusion and fit a series of hierarchical regression models using as predictors a set of key putative determinants of turnover rate, including environmental variability, local species richness and the size of the wider species pool. We represent total anthropogenic impacts on each transect using a synthetic proxy of cumulative effect, the highly spatially resolved Human Modification Index (Kennedy et al. 2019) that combines a suite of stressors including population density and land use. Specifically, we test two propositions: (1) if human impacts on turnover are detectable, and (2) whether the human impacts are consistent across turnover metrics.

By synthesising the response of multiple aspects of species turnover we can then explore the underlying macroecological dynamics. If human impacts on temporal β-diversity are largely a case of the replacement of one avifauna with another, we would expect faster short-term community turnover and higher rates of species accumulation in highly modified areas. However, if human impacts have largely manifested as modifications of the intrinsic dynamics of bird communities, we would expect little impact on the species accumulation rate while the impact on short-term turnover could be in either direction.

## Methods

Unless otherwise noted, all analyses were carried out in R version 4.2.0. All data used is publicly available. Analysis scripts, processed data and model fits are available in repositories (see code availability note).

### Bird Data

We collated yearly presence-absence matrices from the North American Breeding Bird Survey (BBS) using the 2020 release covering 1966 -2019 (Pardieck et al. 2020). BBS data are collected annually on one day during the peak of the avian breeding season along thousands of survey routes across North America. Surveys follow a standardised protocol allowing comparison of diversity patterns through time (Sauer et al. 2017). Each route is 40 km long with 50 observation points at approximately 800 m intervals. At each point, observers carry out a 3-min point count during which every bird observed or heard within ∼400 m radius is recorded. Raw data is then checked and collated centrally.

We excluded species that were not identified to species level or are poorly captured by the BBS surveying methodology (i.e. nocturnal, crepuscular, birds of prey, and aquatic species) following others (Dorazio and Royle 2005, Jarzyna and Jetz 2017). Where taxonomic revisions had split or lumped taxa during the survey period, these were grouped together as a single taxon for consistency. In most cases these are allopatric divisions and so are unlikely to notably impact recorded dynamics on individual transects.

We excluded all surveys that did not pass the BBS quality control (e.g. the weather was poor). To further reduce the impact of unusual weather or poor surveying, if a particular survey had a reported species richness that deviated from the mean richness by more than twice the standard deviation of richness observations for that transect, that survey was excluded (see example in Fig 2, this step excluded 3274 out of 85460 surveys (3.83%)). We retained transects that had associated GPS data for the route (the U.S. Geological Survey Patuxent Wildlife Research Center, http://purl.stanford.edu/vy474dv5024), had more than 11 surveys (in order to generate more than 10 similarity values), and were part of a Bird Conservation Region (BCR, Sauer et al. (2003)) that had at least 25 suitable transects. Our final dataset comprised data from 2692 transects from 27 BCRs, comprised an average of 31.7 (SD = 11.2) surveys, spanning an average of 39.4 years (SD = 11.5). See SI 1 for further summary statistics.

**Fig. 2.**
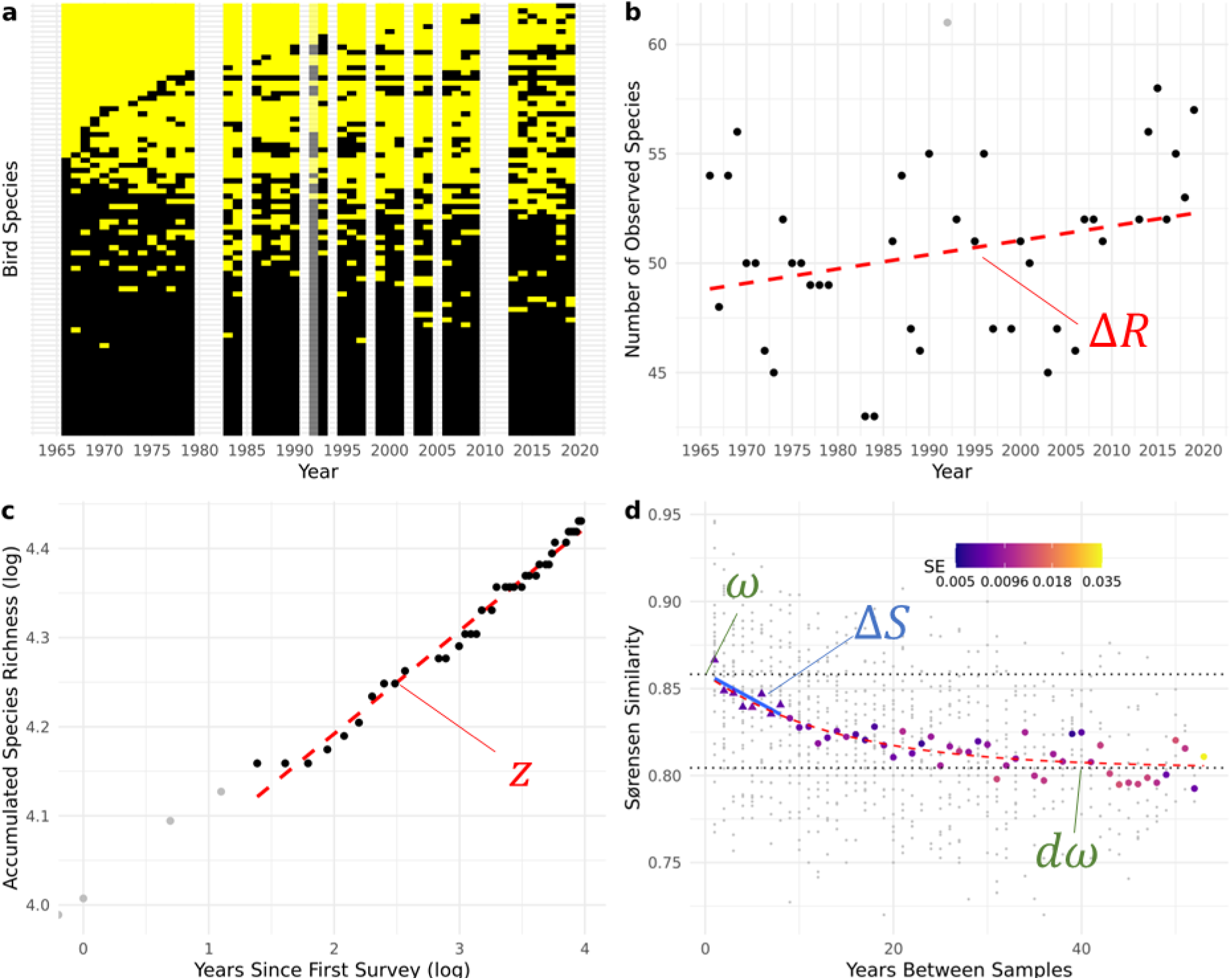
Example transect data illustrating the form of the data and the derived statistics. The example here is the ‘Pine Grove’ transect (C_840_S_02_R_008), located in the ‘Appalachian Mountains’ Bird Conservation Region (Lat: 34.0, Long: -85.7). a) *Species presence-absence matrix*. Only years where there was a survey that passed the BBS quality control are included. Black filled tiles indicate species presence. Species are ordered by the year they are first seen. Data for 1992 is shown in grey – it was excluded from further calculation as the number of species observed was larger than 2 SD of the mean. b) *Observed species richness trend.* Linear model summarising trend is shown with dashed line (slope Δ*R* = +0.0651 sp year^-1^). Excluded year (1992) is shown in grey. c) *Rate of species accumulation* (species-time relationship). Here the total number of species observed at least once since the start of the survey is regressed against time since the first survey. A linear model is fit on a log-log scale, excluding the first 4 points (shown in grey). Here, *z* the slope (exponent) was 0.115. d) *Community similarity decay curve.* Small grey dots show raw Sørensen similarities of each year-pair. Coloured shapes show mean similarity values for each year-difference, with the colour indicating the standard error of the mean used in the model fitting. Fitted initial rate of similarity decay (Δ*S*) for the first 8 points (triangles) is shown with a solid blue line. Note that in order to make a more positive value correspond to a faster turnover, the sign of the slope is reversed to calculate Δ*S*. The similarity decay curve fit (red dashed line) follows *S̅* = ω[(1 − d)*exp*(−Lt) + d], where here d: 0.937, *ω*: 0.858, L: 0.0715. Black dotted lines show fitted maximum observed similarity at t=0, (*ω*), and asymptotic observed similarity (*d* × *ω*).

By studying changes in presence-absence rather than abundance (Shimadzu et al. 2015, Legendre 2019), our study directs focus towards the dynamics of rarer species. Abundance-based analyses frequently find that trends are dominated by a handful of species (Leung et al. 2020, Gotelli et al. 2022), including with this dataset (Di Cecco and Hurlbert 2022). In all transects there is a large group of species that are observed in each year in the transect (SI Fig1.A.g).

### Species Richness Trends and Species Accumulation Rates

We identified trends in the species richness (Δ*R*) by fitting a linear model of species richness against year for each transect (Fig 2b, observed richness were all well above zero justifying this direct approach). We determined the slope of the species accumulation curve (species-time relationship, STR) for each transect, by fitting linear models of *log*(*sp*)∼*z log*(*t*) + *c* (Fig 2c). We based our accumulation curves from the first available year (i.e. a nested design, Carey et al. 2007). To minimise the dominant influence of observation error on the initial rate of species accumulation (White 2004), we excluded the first four samples from the slope fitting. In line with other results (Rosenzweig 1995), these *z* values were highly correlated with exponents derived from the alternative model form *sp*∼*z log*(*t*) + *c*.

### Building Similarity-decay Curves

For each pair of years within each transect, we calculated the Sørensen similarity 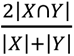 where X and Y are the set of species observed at each sample. Sørensen’s similarity has high capacity to detect change (Santini et al. 2017). However, the preponderance of short-time interval data compared to long-interval data poses a challenge for identifying the statistical confidence in trends at long and short intervals simultaneously (Russell et al. 1995, Collins et al. 2000). For each transect (*i*) and time difference (*t*), we calculated a mean average similarity 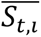 and its standard error to propagate forward into the later stages. Where a particular time difference had only a single representative (making direct calculation of standard error impossible) it was assigned the mean standard error for that transect.

The effective multiple re-use and temporal non-independence of observations from each transect poses a fundamental problem of pseudoreplication. In particular, estimating the value of the average similarity of samples separated by a particular number of years, contributes in a complex way to the overall uncertainty. Because each year of observation potentially contributes to multiple year-pairs, and each year-pair contributing to each estimate of 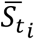 is not independent of each other, our method somewhat underestimates the uncertainty with which the core curve parameters can be estimated.

To calculate the initial rate of similarity decline (Δ*S*) for each transect, we fit a weighted linear model through the 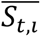 values, up to t = 8 years (Fig 2d). We then use the negative of the slope as a simple measure of turnover, so that higher Δ*S* is a faster decline, for sign consistency with the other metrics of turnover.

### Predictor Data

We generated a set of five predictor variables to use in our regression models (Fig 3). We calculated an ‘observed alpha diversity’ predictor (AL), as the average species richness recorded across the surveys on the transect. The duration of data available (difference in years between the first and last survey at a transect) was included as a year span (YS) moderating variable. We derived a measure of the avian gamma diversity (GM), a proxy of inter-annual environmental variability (CV) and a proxy for human impact (HM) from cross-referencing the transect GPS routes with external data sources. For the fitting process and to aid comparison, AL, GM and YS were scaled to approximately the 0-1 range by dividing by 100, 150 and 50 respectively.

**Fig 3.**
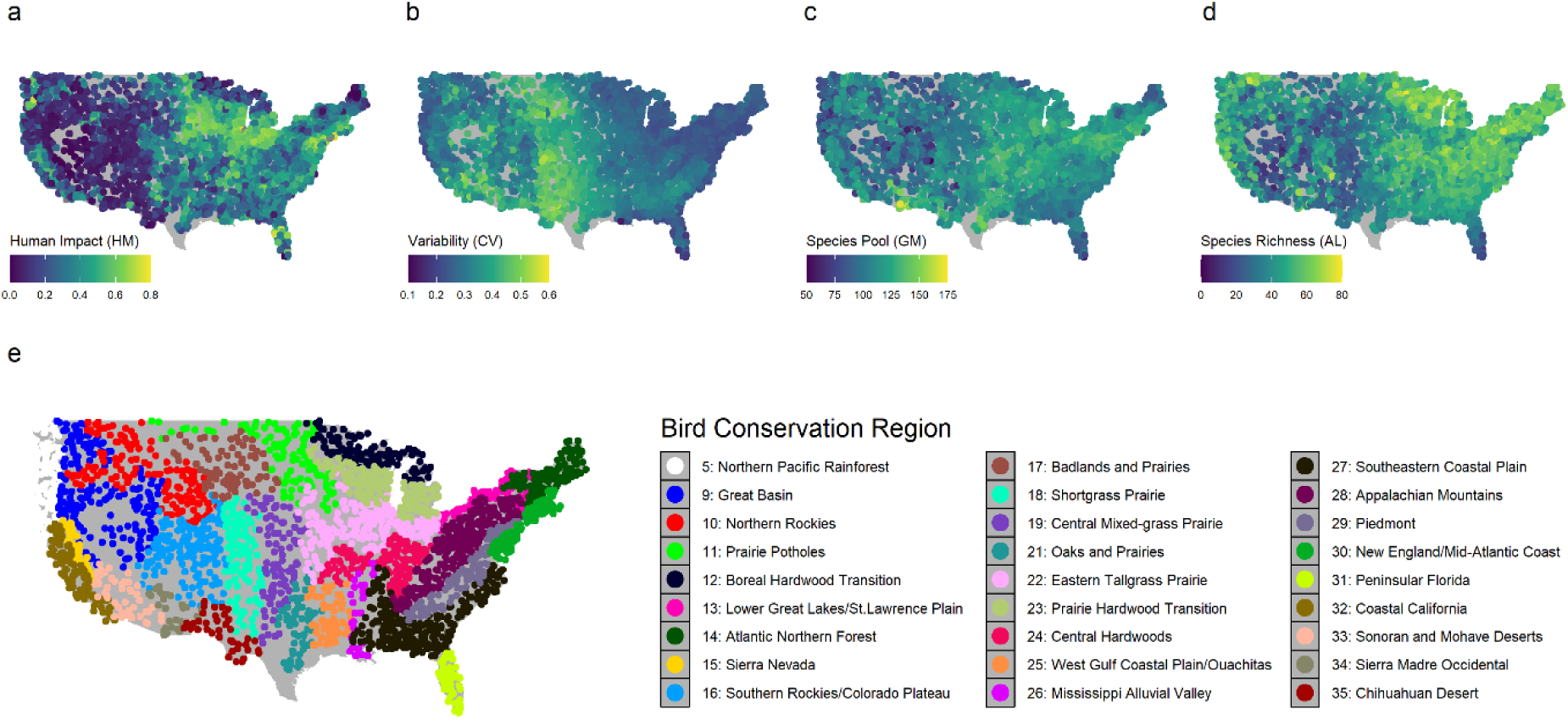
Distribution of the values for four predictor variables for each transect across the United States and allocation of the transects to Bird Conservation Regions (BCRs) of the focal transects. a) Human impact proxy, derived from the Human Modification Index map, b) an environmental variability proxy derived from the coefficient of variation of satellite derived Gross Primary Productivity, c) the size of the species pool, derived from eBird range map data, and d) average observed species richness from the transect surveys. The fifth predictor, the year span of surveys available for each transect is not shown as it shows no meaningful spatial pattern. Points are located at the start of each 40 km route. While predictor variables are strongly correlated at the continental scale, within BCRs the correlations are markedly weaker (SI1). e) the location and description of each Bird Conservation Region used.

To estimate the size of the species pool for our gamma diversity predictor GM we downloaded summer predicted range maps derived from eBird data (November 2021 release, Fink et al. 2020, 2021). These use large-scale citizen science data and finely resolved environmental predictors to generate high-resolution range maps. We used the spatially smoothed (‘medium resolution’, 8.89 km^2^) maps of all species with range map data that were also included in our BBS transects data (375 species). For each transect, we calculated the total number of species ranges that the transect route intersected. Hence, GM is also partly a measure of the heterogeneity of habitats the route crosses. For our measure of inter-annual variability CV, we used high-resolution MODIS satellite derived Gross Primary Productivity (GPP) data (MOD17A2H V6) accessed through Google Earth Engine (https://developers.google.com/earth-engine/datasets/catalog/MODIS_006_MOD17A2H). For a 1km buffer around each transect route, we calculated the mean GPP for each year from 2001 to 2021 (the period for which complete and consistent data was available). We then calculated the coefficient of variation of annual productivity for each transect. To reduce the effect of high-outliers, we square-root transformed the coefficient of variation to generate our variability proxy ‘CV’.

For our measure of anthropogenic impact HM, we used a global human modification raster map at 1km scale (Kennedy et al. 2019), which combines 13 anthropogenic stressors (including population density, agricultural land use, transport infrastructure, fragmentation and night-time lights) into a single 0-1 ‘Human Modification Index’. We intersected the transect routes with this map, and took a mean value along the route.

### Fitting Similarity Decline Curves and Impact of Predictors

The response of the species richness trends (Δ*R*) and rates of species accumulation (*z*) to our suite of predictors was tested using linear multiple regression models, fit separately within each BCR.

We fit a curve function through the decline in similarity values through time (Fig 2d) with a Bayesian hierarchical model for each BCR separately using the Hamiltonian Monte Carlo sampler STAN (Carpenter et al. 2017) accessed through the *brms* (version 2.18.0, Bürkner 2018) interface in R. Response variable uncertainty in each 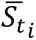 estimate was incorporated by the estimated standard error of the point as calculated above. The core model for each transect was a bounded exponential decay model of the form:

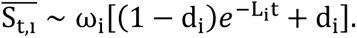

Here, *L* represents the initial similarity decay rate and *d* the asymptotic community similarity. These are scaled by a *ω* term that represents observation repeatability. At a time-difference of 0 there should be complete similarity between observed communities with consistent detection. The *ω* term captures the proportional reduction in observed similarity between samples attributable to measurement error – for example species that were present in both samples but unobserved in one, or alternatively transient species that were observed once but not truly resident. It is fit at the level of the survey route, capturing both the difficulty in observing birds on the route and the varying skill of observers assigned to the route.

Note that a levelling off in turnover is only clearly visible in a subset of similarity decay curves (See Figure S4). However, although in many cases the observed similarity decay cannot be clearly distinguished from a linear decline via classic model comparison (Penny et al 2022), ultimately similarity is bounded 0-1. To handle this while avoiding imposing a high threshold, we assume an exponential decay model but fully propagate the uncertainty in the model coefficients. Hence, while some time series contain only the minimal information about the upper limit of where levelling off will take place, these values will have a high associated uncertainty and as such will contribute substantially less to the final results.

Each parameter is in turn described by the predictors with linear models:

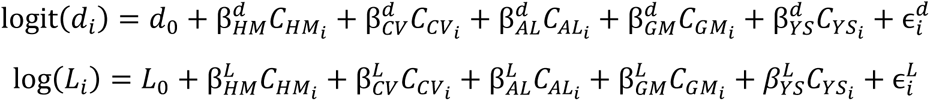

and

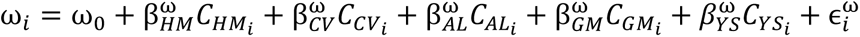

The transect-level hierarchical terms 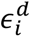, 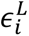 and 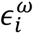 are considered to be drawn from Gaussian distributions with fitted standard deviations. Here β terms represent coefficients and *C* terms the centred vectors of predictor data. The residual error term is also allowed to vary between transects, again following a Gaussian distribution. Priors were chosen to be relatively weak with the aim to aid convergence without impacting the final result. See SI 6 for full model details and for details of the priors assumed. For each BCR, predictors were first centred by subtracting the mean for that BCR. For each model four chains each of total length 10’000 were run, discarding the first 5000 as burn-in. Key convergence statistics indicated convergence was achieved. Raw STAN code and model fits are available in the online repository. Separately, to estimate the best fit values of the curve parameters we repeated the fitting process but without the regression terms. All other aspects were held the same.

### Cross-BCR Analysis

Finally, for each of the six responses (Δ*S*, Δ*R*, *z*, ω, *d*, *L*) we conducted a simple meta-analysis across all BCRs to identify if a consistent effect of each predictor can be identified, where each BCR included in our analysis was treated as a separate ‘study’. For this we used a linear model fit with *brms* to estimate the mean effect (coefficient) of each predictor across the BCRs, incorporating the standard error of the coefficient estimates from the within-BCR model fits.

## Results

The total distribution of fitted response parameters and impact of predictor variables across all regions are shown in Fig 4. Across the 27 focal BCRs, there was a general consistency in the impact (or lack of impact) of our putative determinants of the similarity decay curve (SI 2). Overall, higher human modification was associated with greater observation repeatability (*ω*, Fig 4a), lower asymptotic similarity (*d*, Fig 4b) and slower rate of similarity decay (*L*, Fig 4c). Higher average observed species richness were associated with significantly greater *ω*, lower L and greater d. Larger regional species pools were associated with marginally lower *ω*, but no clear impact on *d* or *L*. Environmental stochasticity, as described by productivity variability, was not consistently influential and study duration had negligible impact on the parameters of the similarity decay curve.

**Fig.4.**
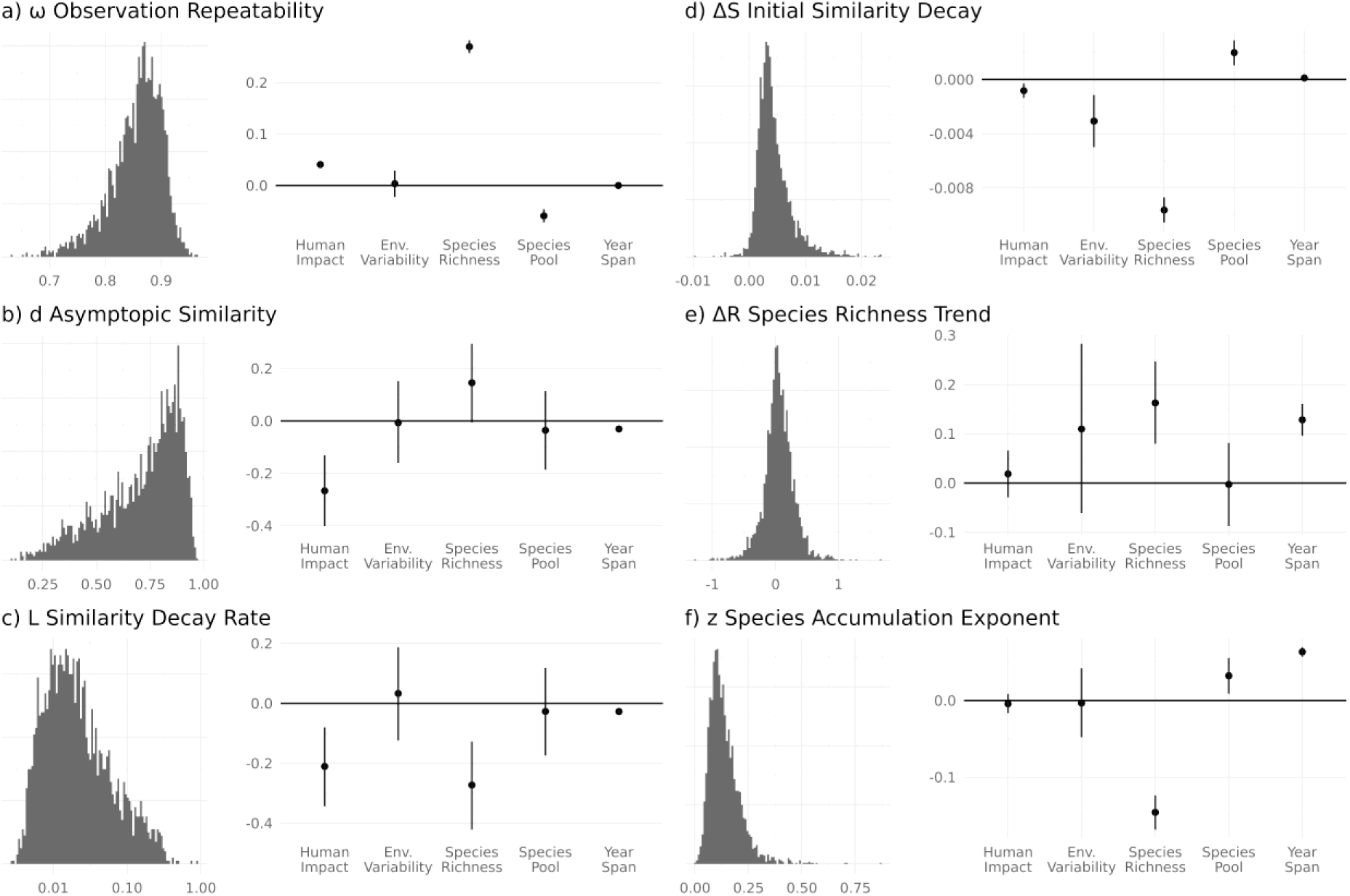
Histograms of fitted turnover statistics across all transects and estimated effects of the key predictor variables across all Bird Conservation Regions (BCRs). The overall impact of the predictors shown on each right column is effectively a meta-analysis of the coefficients identified in each of the 27 focal BCRs. Separate BCR-level results are shown in Figure S2. Predictor variables are each scaled to approximately 0-1 range prior to centring (Year Span /50, Species richness /100, Species pool /150), but the magnitude of impact should not be considered directly comparable between predictors. Lines show central 90% of posterior distribution.

The initial rate of similarity decay (Δ*S*), as measured by a direct linear model through the early stages of the similarity decay curve, was slightly reduced by human modification (Fig.4d). This implies higher levels of anthropogenic impact were associated with slower short-term turnover. However, initial rates of decay were much more influenced by the observed species richness (higher average observed species richness was strongly associated with slower initial decline in similarity) and the impacts were comparatively inconsistent across BCRs (SI 2).

Observed species richness was largely consistent through time on most transects (Fig 4e histogram), in line with previous results with this dataset (Schipper et al. 2016, although see Jarzyna and Jetz 2017). Across all transects, the mean change in richness was low: +0.053 species per year, equivalent to around two additional species between the start and the end of the survey period. With an average of 45.7 species seen on each transect, this is a small proportional change. This near-balance between gains and losses justifies our focus on a simple measure of community similarity rather than seeking to partition out the dissimilarity due to changing species richness.

The accumulation of observed species after *t* years (*sp*_*t*_) could be well described by a standard STR function: log(*sp*_*t*_) ∼z log(*t*) + c where the overall mean species accumulation exponent (*z*) across all transects was 0.156 (Fig 4f). Average observed species richness was consistently negatively correlated with this exponent, in line with previous analyses (White et al. 2006). Transects with a longer duration of data had marginally higher exponents. The Human Modification Index had no detectable impact, while other relationships with the predictor suite were small in relative magnitude and inconsistent between BCRs (SI 2).

## Discussion

We found pervasive human impacts on species turnover between the bird communities across habitats, but crucially these effects depended on the aspect of ‘turnover’ considered. Taken together they give a deeper picture of the fundamental ecological dynamics with significant implications for the interpretation of short-scale monitoring programs for conservation objectives. Our proxy for anthropogenic disruption, the Human Modification Index (Kennedy et al. 2019), was associated with lower asymptotic similarity (*d*). By contrast, the exponent of the species-accumulation curves (*z*) was not clearly related to the Human Modification Index, once differences in average species richness had been accounted for. Furthermore, the initial short-term rate of similarity decline (ΔS), a frequently examined method of assessing turnover, showed marginally *slower* turnover in areas of higher human modification. Hence, core measures of ‘turnover’ appear to give contrasting inferences of anthropogenic impact.

These differences have significant implications for the use of single measures of observed turnover for conservation objectives – inferences of one aspect of turnover may not translate to others. A frequent objective of turnover studies has been to determine a single annualised rate of species extinction and colonisation (Diamond 1969). Multiple authors have sought to identify ideal sampling frequencies and analyses to better capture this quantity (Diamond and May 1977, Nichols et al. 1998, Collins et al. 2000, Ontiveros et al. 2021), traversing the twin perceived threats of missing species extirpation and recolonisation between censuses (crypto-turnover) and of failing to consistently observe present species (pseudo-turnover). Our results suggest that this may risk obscuring important dynamics and suggests that arguments identifying higher short-term turnover as an indicator of anthropogenic impact and hence conservation focus (Blowes et al. 2019) may need to be reassessed.

The contrasts also offer the opportunity to qualitatively triangulate the relative role of intrinsic and extrinsic processes in determining how anthropogenic effects translate into biodiversity changes (Fig. 1). Identifying the extent of intrinsic turnover is crucial to understanding whether observed changes in species composition are a concern or in fact an indication of a healthy resilient ecosystem undergoing ‘normal’ dynamics (Magurran 2016). Short-term turnover rates in themsevles does not have enough information to distinguish intrinsic and extrinsic turnover. Observations of definitive levelling off of similarity over long time scales seen in some transects (SI Figure 4b) suggests that, in these transects at least, intrinsic turnover dominates the observed turnover dynamics since sustained and directional extrinsically driven turnover would be expected to drive continually declining similarity. However, for the majority of transects where levelling off is not clearly observable further information is needed.

The rate of species accumulation over sequential samples, often referred to as the species-time relationship (‘STR’, White 2004), provides a route to qualitatively differentiate the impact of humans on intrinsic and extrinsic processes. Since extrinsic changes can be expected to result in the arrival of ‘new’ species at a faster rate compared to intrinsic turnover, increases in extrinsically driven turnover would be expected to increase the exponent *z*. The minimal impact of humans on the rate of species accumulation suggests that, for this turnover measure at least, any human-driven increases in the arrival of novel species are being matched by a reduction in the background rate of turnover. Changes in the environment due to human modification are impacting long-term turnover through shifts in the status of species from ‘rarely seen’ to ‘frequently seen’ (and vice-versa) rather than simply an increased rate of appearance of ‘new’ species.

To summarise, at the transect scale of our study human impacts are generally causing a slower, but deeper, mixing of the species pool that is largely already present in the wider metacommunity. While the relative balance between the two can’t be precisely quantified, intrinsic dynamics appear to have a key role in determining the rate of observed turnover, upon which are overlaid the impacts of extrinsic drivers of turnover, such as land use change (Rittenhouse et al. 2012, Di Cecco and Hurlbert 2022). Concordant results were also found in long term surveys of Canadian trees (Brice et al. 2019) where, although community wide functional trait shifts driven could be detected, the role of ongoing disturbances had a greater impact on turnover than environmental change.

A moderate slowdown of short-term turnover due to human impacts may appear counter-intuitive, particularly as known reductions in the population abundance of many species (Schipper et al. 2016, Rosenberg et al. 2019) may erode the sustainability of local species populations. However, there are several possible explanatory mechanisms for this effect. Any possible impact pathways need to be considered in the light of the observation of only slight trends in species richness across the study period (Schipper et al. 2016). This implies that overall extirpation and colonisation on the transects are approximately balanced. Most directly, observed slow-downs of turnover with human impact could be a manifestation of a shifting baseline – if more highly impacted areas had already lost many rare and ephemeral species before the start of the survey period, this could explain the reduction in short-term turnover. This hypothesis is hard to directly test without older baseline data, however, we note that there was minimal correlation between observed species richness and our human modification proxy (within BCR average Pearson’s correlation = -0.0006, SI 1bii).

Human modification could also be associated with slower turnover directly, by reducing the causes of intrinsic turnover, in particular by slowing colonisation rates or weakening interspecific interactions. However, precisely diagnosing specific mechanisms from observation data such as this is a considerable challenge. The effective balance in species richness could be attributable to extinctions contributing to opening up niches for new colonists to fill, or colonisation may lead to exclusion of previous residents from the community. Directly distinguishing gap exploitation from displacement driven turnover is not straightforward from observational data, particularly as individual cases of extirpation and colonisation can lead to either homogenisation or heterogenization between two samples (Olden and Rooney 2006). In a scenario corresponding to a saturated (structurally unstable, Rossberg et al. 2017) community, if human impacts led to a reduction in the rate of colonisation of sites along the transect from the wider metacommunity the short term rate of turnover would fall.

At the scale of presence/absence on BBS transects we are unaware of direct evidence for widespread competitive exclusion between species, although there are examples of competitive exclusion in birds in other systems (Martin and Bonier 2018, Freeman et al. 2022). However, given the complexity of possible interactions, we would not necessarily expect clearly observable patterns of exclusion to result from competition (Rossberg 2013, Blanchet et al. 2020). Turnover driven by continual immigration events has been shown to be able to reproduce a host of observed patterns in models where the total diversity of a community is in some way limited (O’Sullivan et al. 2019), such that each colonisation tends to lead on average to an extinction from the community. Colonisation pressure is likely to be positively linked to the diversity of the species pool (i.e. gamma diversity) and we do see small positive associations between the species pool and both the initial similarity decline and species accumulation exponent. Further research into the underlying mechanisms behind intrinsic turnover will be highly valuable. Analysis of trends in functional diversity (Jarzyna and Jetz 2017) may be able to identify signals of trait-based species sorting.

The four ecological predictors that we built into our regression models each capture a variety of processes and need to be interpreted with care. Human impacts have not been consistent through time, and the state of modification at the end of the time period (the Human Modification Index is based on data with a median year of 2016), will obscure differences between areas where there was large scale land-conversion during the last 50 years, and those where human impacts were already at a high level long before the surveys started. Further, the BBS survey was not initially designed in a fully structured way, and as such there are potentially problematic links between survey period and ecological predictors. For example, earlier-starting transects with longer data spans tend to be in relatively more human-modified areas, while recent heavy land use change has been found to be associated with the early stopping of transect data collection (Zhang et al. 2021). As a quantitative assessment of the total extent of changes to turnover across the US may be beyond reach we therefore focus on identifying the direction of links between our predictors and biodiversity responses.

While all the transects cover the same distance and use the same protocol, there are significant differences in their route shape and the heterogeneity of the habitat that they cross. This heterogeneity is known to influence observed species richness (Farwell et al. 2020). Our measure of average observed species richness is therefore unlikely to represent only the classic understanding of alpha diversity as being the number of species existing in a particular point location. A significant habitat heterogeneity component could provide a mechanistic explanation of why the initial rate of similarity decay is slower with higher observed species richness: increased habitat heterogeneity could reduce the number of potential colonists in neighbouring areas able to drive intrinsic turnover (O’Sullivan et al. 2021). Similarly, our metric of the size of the species pool (the number of bird species ranges the transect route crosses), will also be affected by the heterogeneity of habitat crossed by the route. This may explain the negative relationship between species pool and observation repeatability. A route that crosses a greater variety of habitat subtypes will inevitably have less observation time in each, and a higher likelihood of missing specialists. Although the range maps are fundamentally porous and are so likely to overestimate the real species pool at any given location (Hurlbert and White 2005), it is reasonable to expect a good correlation.

Our intercept term, ω, is designed to account for the apparent dissimilarity in repeat samples attributable to undetected resident species or the observation of vagrant species passing through, but not truly resident. Our framework assumes that detectability was approximately constant through time. The nearly constant observed species richness suggests this assumption is reasonable. With perfect surveys, similarity at t=0 should be 1. With the bulk of ω values falling between 0.8 and 0.9, missed species contribute notably to apparent turnover between surveys. Previous studies that have directly assessed measurement error using raw stop-level count data have identified strong differences between transects in average detectability of species (Boulinier et al. 1998, Jarzyna and Jetz 2016). It is likely that differences in detectability between transects are behind the strong positive association between observed species richness and ω. Our metric of human impact was also associated with increased ω. This could be a measurement effect, such as greater habitat openness and hence fundamental detectability, or a biological effect, where humans cause extirpation of rare, hard to detect species (and consequently raising the average detectability of the remainder).

## Conclusion

By taking a holistic view of different aspects of species turnover (Magurran et al. 2019), we have been able to identify a pervasive anthropogenic impact on ecological community dynamics. Highly human impacted regions may not be showing notable changes in species richness, and in fact show somewhat slower rates of short-term community turnover, but they appear to show deeper long-term community change through time as the community undergoes fundamental changes in structure. This effect appears to be largely driven by changes in the patterns of temporal occupancy within the bird community, rather than the arrival of ‘new’ species. That said, the recent past is not necessarily a guide to the future, and rates of extrinsic change (particularly climate impacts) are likely to accelerate in the coming decades (Doak and Morris 2010, Nolan et al. 2018). By focussing on a single taxonomic group with largely consistent sampling protocols, we have been able to undertake a more detailed analysis, but results from meta-analyses with broader scope (Antão et al. 2020, Pilotto et al. 2020) suggest that we should expect considerable differences between taxa and biomes. Our results highlight the pressing need to understand mechanistically the turnover in ecological communities in order to better interpret future changes.

## Acknowledgements

We gratefully acknowledge and thank the thousands of BBS survey participants who annually perform and coordinate the survey. This research. We thank Anne Magurran, Simon Mills, Tom Brewer, Jacob O’Sullivan and Emmanuel Nwankwo for their comments on manuscript drafts. This work was funded by NERC grant NE/T003510/1 to AGR. JCDT is also funded by a Leverhulme Trust Early Career Fellowship (ECF-2022-666) and used utilised Queen Mary’s Apocrita HPC facility.

## Author Contributions

JCDT conceptualised, designed and conducted the research and wrote the manuscript. AGR contributed to the analysis and results interpretation. Both contributed to manuscript editing.

## Supplementary Information

**Figure S1.**
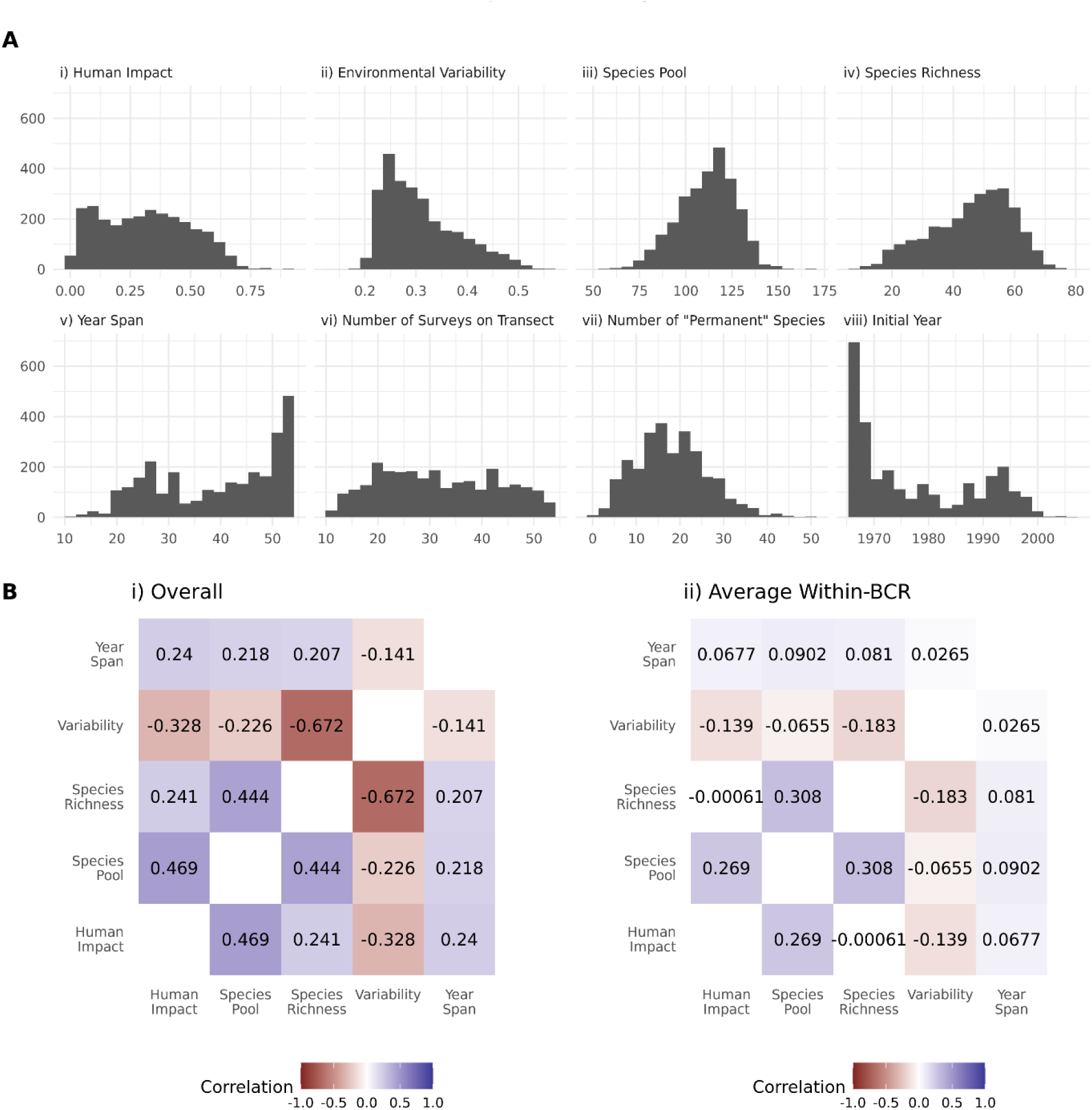
Predictor variable data summaries. A) Distribution of key predictors and summary statistics. Histograms are shown across all transects and values are shown before scaling used in regressions. Permanent species (g) are operationally defined as those seen in every year the transect survey was included. B) Correlation between key predictor variables i) across all transects ii) within each BCR, then averaged.

**Figure S2.**
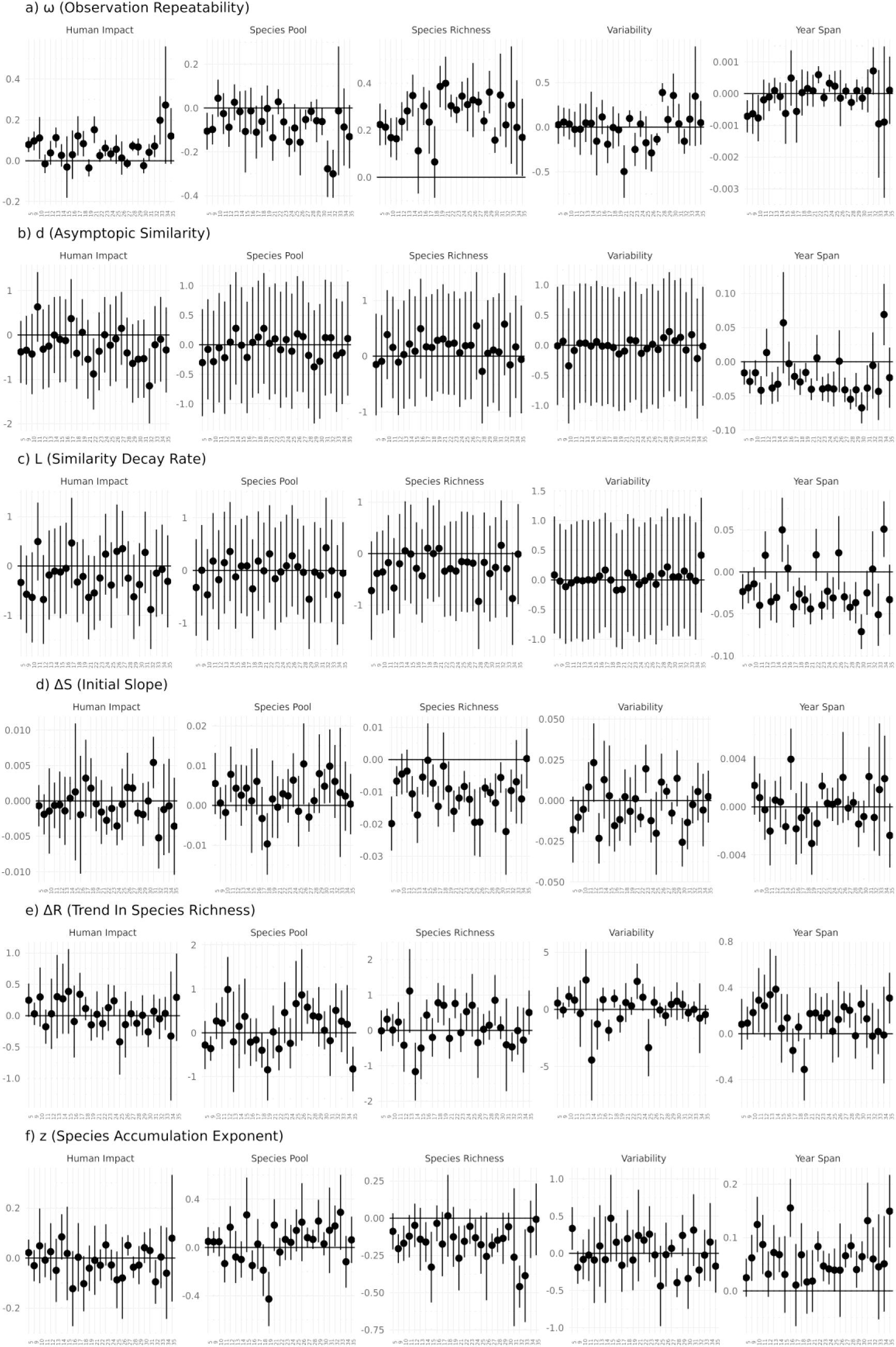
Fitted coefficients for each predictor for each Bird Conservation Region (BCR). Each BCR is given by its number (see Fig 2 for description). Points show the mean estimate, while error bars show two standard errors around each mean estimate.

**Figure S3.**
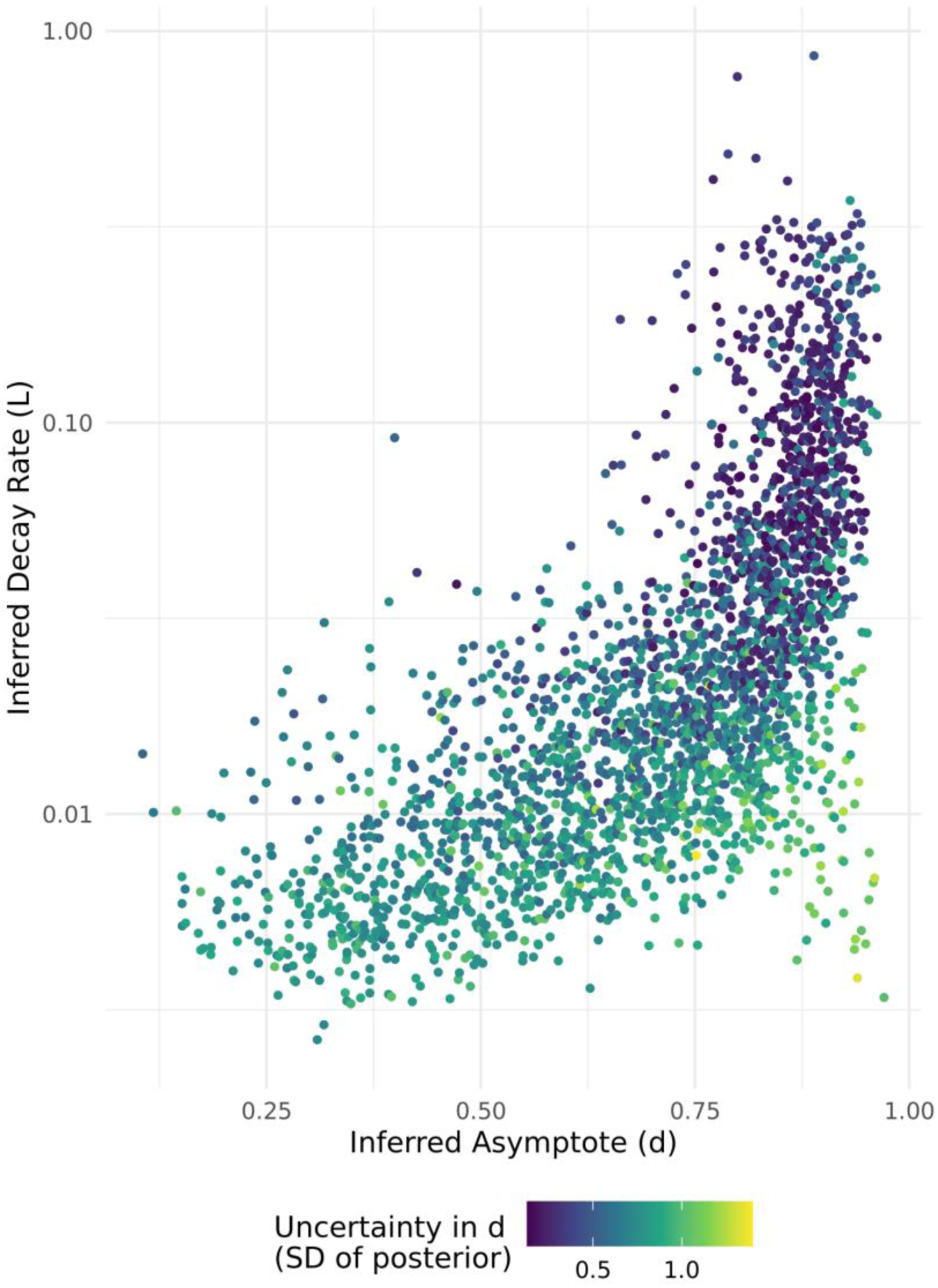
Correlation in estimated curve parameters. Posterior means of similarity decay curve parameters when directly fitted i.e. without the predictor variables. Results are shown across all Bird Conservation Regions. Points are coloured by the uncertainty in the asymptote (standard deviation of the posterior on the logit scale). There is a clear positive correlation between the d and L parameters from which we can broadly identify three archetypes of observable temporal similarity dynamics. At the top right, ‘fast’ transects with high L and high d, show a quick decline to a clear high asymptote. At the bottom right, ‘slow’ transects with low L and low d show a clear saturation, but take considerably longer to reach that point. Thirdly, in the bottom right there are a smaller group of ‘uncertain’ transects where the asymptote is uncertain. These are frequently transects with shorter available time series (Year Span).

**Figure S4.**
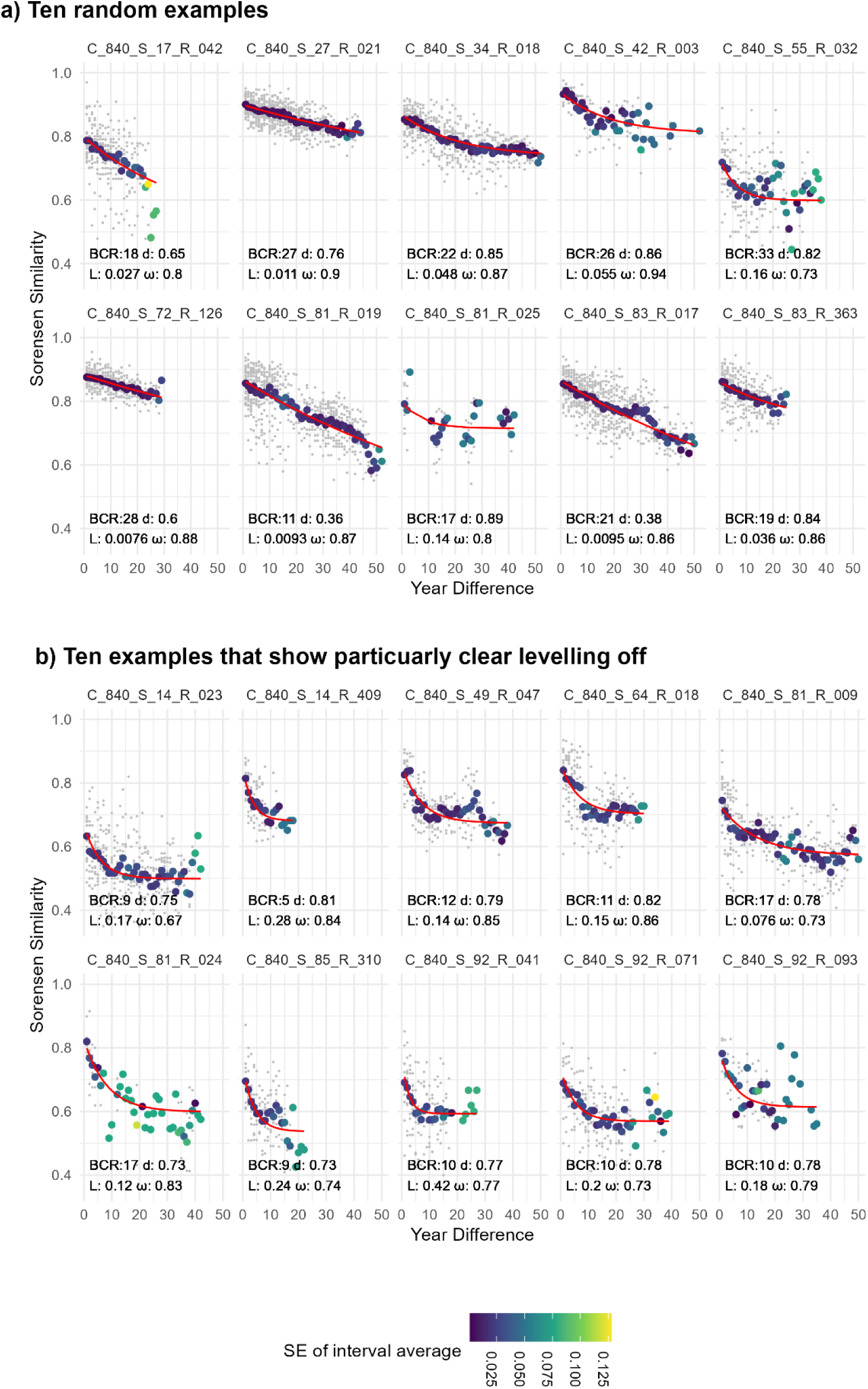
Randomly selected examples of similarity-decay curves. Colour of each point shows the confidence assigned to each estimate of the mean similarity for that year difference, calculated as a standard error. Small grey dots are each similarity value underlying these averages. Red line shows the directly fit (i.e without the predictor variables, but with the partial pooling) decay curve using the mean of the posterior values. Transects are titled by country code (US = 840), state code and route number. Curve parameters and BCR number are shown for each transect.

### SI 5

Details of hierarchical model used for fitting similarity decay curve. The model is described in the main text methods, but here we give further specific details. Each BCR was fit separately in *brms* 2.18, an R package that calls STAN. All code (including the generated STAN code) is available online, as well as the fitted model objects. The central *brms* call was:

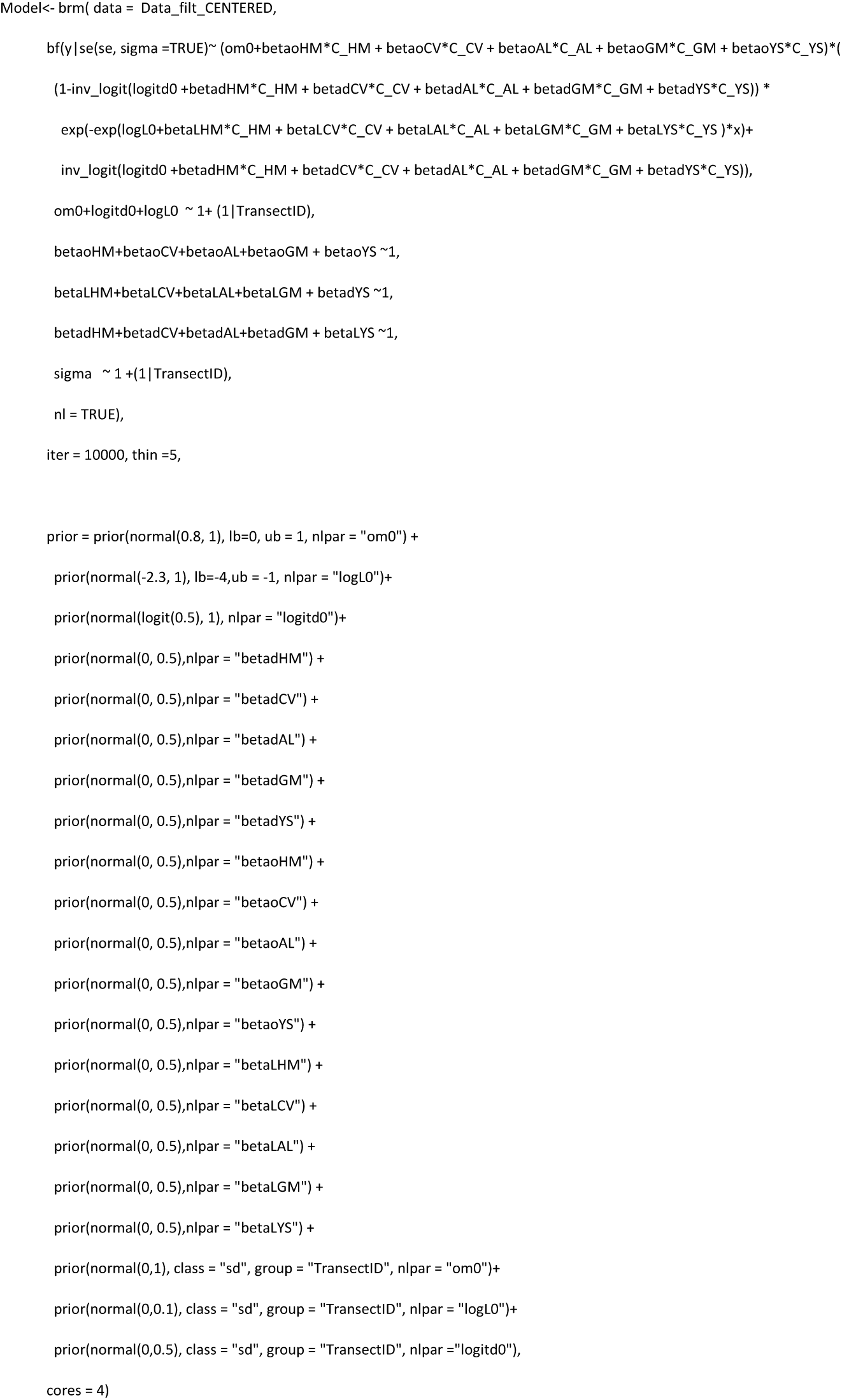

where the input data is separate for each BCR and has columns:

- x = Difference in years between samples
- y= Average Sorensen similarity
- se = Standard error of the estimate of y.
- TransectID = character string identifying the transect
- C_HM = Human modification Index value of transect (centred)
- C_CV = Environmental Variability value of transect (centred)
- C_AL = Observed Species Richness value of transect(centred)
- C_GM = Species Pool value of transect (centred)
- C_YS = Year Span value of transect (centred)

The full model to determine the likelihood (excluding the priors) of each average Sorensen similarity 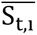 with time-difference *t* on transect *i* is:

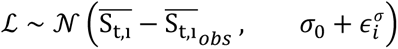

where

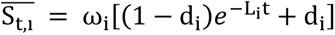

and:

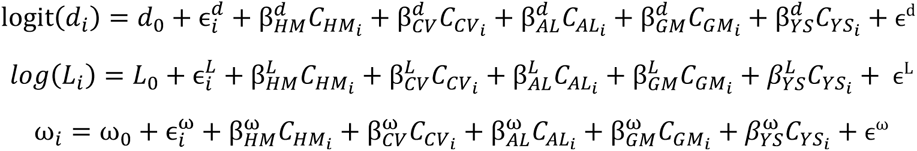

with the hierarchical pooling terms to specify different intercepts for term for each transect *i* are defined based on Normal distributions, each with a fitted standard deviation:

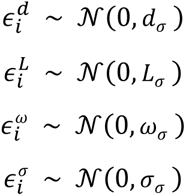

Priors not manually specified above follow the weak defaults of brms. The complete list of priors were:

|  |  | Distribution | Parameters | Bounds |  | Notes |
| --- | --- | --- | --- | --- | --- | --- |
|  |  |  |  | Lower | Upper |  |
| Regression intercept terms |  |  |  |  |  |  |
| | $d_0$ | Normal | Mean =0, sd =1 | | | NB logit term |
| | $L_0$ | Normal | Mean = -2.3, sd = 1 | -4 | -1 | NB log term |
| | $\omega_0$ | Normal | Mean = 0.8, sd = 1 | 0 | 1 | |
| Regression Main Effect Terms |  |  |  |  |  |  |
| | All $\beta$ | Normal | Mean = 0, sd = 0.5 | | | |
| Regression Error Terms |  |  |  |  |  |  |
| | $\epsilon^d$ | Student t | Df = 3, loc = 0, scale = 2.5 | 0 | | |
| | $\epsilon^L$ | Student t | Df = 3, loc = 0, scale = 2.5 | 0 | | |
| | $\epsilon^\omega$ | Student t | Df = 3, loc = 0, scale = 2.5 | 0 | | |
| Partial pooling across transects |  |  |  |  |  |  |
| | $d_\sigma$ | | Mean = 0, sd = 0.5 | 0 | | |
| | $L_{\sigma}$ | | Mean = 0, sd = 0.1 | 0 | | Prior on difference between transects |
| | $\omega_{\sigma}$ | | Mean = 0, sd = 1 | 0 | | |
| Overall Error |  |  |  |  |  |  |
| | $\sigma_0$ | Student t | Df = 3, loc = 0, scale = 2.5 | | | Average error between estimate and observation |
| | $\sigma_{\sigma}$ | Student t | Df = 3, loc = 0, scale = 2.5 | 0 | | Difference between extent of error between transects |

